# Variability in error-based and reward-based human motor learning is associated with entorhinal volume

**DOI:** 10.1101/2020.05.27.119529

**Authors:** Anouk J. de Brouwer, Mohammad R. Rashid, J. Randall Flanagan, Jordan Poppenk, Jason P. Gallivan

## Abstract

Error-based and reward-based processes are critical for motor learning, and are thought to be mediated via distinct neural pathways. However, recent behavioral work in humans suggests that both learning processes are supported by cognitive strategies and that these contribute to individual differences in motor learning ability. While it has been speculated that medial temporal lobe regions may support this strategic component to learning, direct evidence is lacking. Here we first show that faster and more complete learning during error-based visuomotor adaptation is associated with better learning during reward-based shaping of reaching movements. This result suggests that strategic processes, linked to faster and better learning, drive individual differences in both error-based and reward-based motor learning. We then show that right entorhinal cortex volume was larger in good learning individuals—classified across both motor learning tasks—compared to their poorer learning counterparts. This suggests that strategic processes underlying both error- and reward-based learning are linked to neuroanatomical differences in entorhinal cortex.

**Significance Statement:** While it is widely appreciated that humans vary greatly in their motor learning abilities, little is known about the processes and neuroanatomical bases that underlie these differences. Here, using a data-driven approach, we show that individual variability in error-based and reward-based motor learning is tightly linked, and related to the use of cognitive strategies. We further show that structural differences in entorhinal cortex predict this intersubject variability in motor learning, with larger entorhinal volumes being associated with better overall error-based and reward-based learning. Together, these findings provide support for the notion that the ability to recruit strategic processes underlies intersubject variability in both error-based and reward-based learning, which itself may be linked to structural differences in medial temporal regions.

## Introduction

The human brain’s capacity to learn new motor commands is fundamental to almost all activities we engage in. Traditionally, such learning has been viewed as an implicit, procedural process of the motor system, with neural studies focusing on brain areas in the frontoparietal cortex, striatum or cerebellum (Doya, 2000; Lalazar and Vaadia, 2008; Taylor and Ivry, 2014). Only relatively recently have studies demonstrated that cognitive systems, including processes related to strategy use and memory, can bolster or interfere with aspects of motor learning (Mazzoni and Krakauer, 2006; Keisler and Shadmehr, 2010; Taylor and Ivry, 2011; Seidler et al., 2012; Holland et al., 2018). It has been speculated, but not yet shown, that regions in the medial temporal lobe (MTL) may contribute to this cognitive component to motor learning.

In error-based learning, the form of learning by which we refine and adjust our movements to changes in the body or the environment based on observable errors, the use of cognitive strategies (often termed the ‘explicit’ component) has been shown to drive large, rapid changes during early learning (Taylor and Ivry, 2011; Taylor et al., 2014). This is in contrast to the implicit process, which contributes to learning in parallel but in a nonconscious, gradual fashion. Whereas the reliance of the implicit process on the cerebellum is well established (Smith and Shadmehr, 2005; Tseng et al., 2007), the neural basis of the explicit component remains speculative. Evidence from neuroimaging, aging, and lesion studies have implicated areas in the prefrontal cortex in explicit strategies (Shadmehr and Holcomb, 1997; Della-Maggiore and McIntosh, 2005; Taylor and Ivry, 2014). In addition, it has been suggested that regions in the MTL, given their role in declarative processes, may be involved in the explicit component to motor learning (Doyon and Benali, 2005; Taylor and Ivry, 2014; de Brouwer et al., 2018).

In reward-based learning, the form of learning in which motor commands are updated by signals related to success or failure (Sutton and Barto, 2018), the use of cognitive strategies have also been shown to play a pivotal role in performance (Codol et al., 2018; Holland et al., 2018). Conventionally, reward-based learning has been shown to involve neural circuits in the basal ganglia and striatum (Doya, 2000), but there is also some emerging evidence to suggest contributions from MTL regions (Gershman and Daw, 2017; Duncan et al., 2018). A key feature of reward-based learning is that it is achieved through exploration (i.e., the brain figuring out motor commands that increase success). Insofar as such exploration is facilitated by strategies, MTL structures may also contribute to performance during reward-based motor learning.

The role of MTL regions in declarative memory and spatial navigation have been well established (Eichenbaum and Cohen, 2014). In humans, for example, anatomical imaging methods have demonstrated clear links between individual differences in hippocampus and/or entorhinal cortex volume with performance in memory and navigation tasks (Maguire et al., 2000; Rodrigue and Raz, 2004; Whiteman et al., 2016; Sherrill et al., 2018). It is increasingly recognized, however, that the hippocampal-entorhinal system can support more abstract relational representations (Tavares et al., 2015; Constantinescu et al., 2016; Horner et al., 2016; Aronov et al., 2017), and forms a ‘cognitive’ map for representing goals and relating objects and actions within a spatial context (Tolman, 1948; O’Keefe and Nadel, 1978). Such maps are likely to be critical when forming new action-outcome associations, as is the case when searching for and implementing strategies during motor learning.

Here we asked whether individual differences in motor learning performance are linked to hippocampal and entorhinal volume in humans. To examine this, we had human participants undergo a structural neuroimaging session in addition to performing separate error-based and reward-based learning tasks, both known to elicit the use of strategies. We show that learning performance in both motor tasks is directly related and that better overall learning across tasks is associated with larger entorhinal cortex volume.

## Materials and Methods

### Participants

The current study used a subset of participants (N=34; 18 men and 16 women, aged 20-35 years) from a larger cohort study (registered at https://osf.io/y8649) in which 66 right-handed paid volunteers underwent structural and resting state MRI scans. Our thirty-four participants took part in an error-based and reward-based motor learning testing session in addition to participation in the main study. One of these participants was excluded from further analysis because of a high number of invalid trials in the error-based learning task (>25%), thus leaving 33 participants for analysis.

The main experiment and motor learning follow-up tasks were approved by the Queen’s University Health Sciences Research Ethics Board, and participants provided written informed consent before participating in the main experiment and in the motor learning session. The motor learning session took approximately an hour and 45 minutes and participants were compensated $20 for their time. The methods, hypotheses and data analyses for the current study were pre-registered on OSF (https://osf.io/7prq5).

### Neuroimaging

#### Procedure

The day prior to each participants’ MRI scan, participants completed a biofeedback session in a simulated (mock) MRI scanner to become familiar with the MRI environment and to learn to minimize head movement. During the biofeedback session, participants viewed a 45-minute documentary with a live readout trace of their head motion overlaid. When their head motion exceeded an adaptive threshold, the documentary was paused for several seconds while static was played on the screen along with a loud, unpleasant noise. The next day, MRI data were collected over the course of a 1.5-hour session using a 3T whole-body MRI scanner (Magnetom Tim Trio; Siemens Healthcare). We gathered high-resolution whole-brain T1-weighted (repetition time [TR] 2400 ms; echo time [TE] 2.13 ms; flip angle 8°; echo spacing 6.5 ms) and T2-weighted (TR 3200 ms; TE 567 ms; variable flip angle; echo spacing 3.74 ms) anatomical images (in-plane resolution 0.7 ⨉ 0.7 mm_2_; 320 ⨉ 320 matrix; slice thickness 0.7 mm; 256 AC-PC transverse slices; anterior-to-posterior encoding; 2 ⨉ acceleration factor) and an ultra-high resolution T2-weighted volume centred on the medial temporal lobes (resolution 0.5 × 0.5 mm_2_; 384 ⨉ 384 matrix; slice thickness 0.5 mm; 104 transverse slices acquired parallel to the hippocampus long axis; anterior-to-posterior encoding; 2 x acceleration factor; TR 3200 ms; TE 351 ms; variable flip angle; echo spacing 5.12 ms). The whole brain protocols were selected on the basis of protocol optimizations designed by Sortiropoulos and colleagues (2013). The hippocampal protocols were modeled after Chadwick and colleagues (2014). In addition, we acquired two sets (right-left direction and left-right direction) of whole-brain diffusion-weighted volumes (64 directions, b = 1200 s/mm_2_, 93 slices, voxel size = 1.5 ⨉ 1.5 ⨉ 1.5 mm_3_, TR 5.18 s, TE 103.4 ms; 3 times multiband acceleration), plus two extra B0 scans gathered separately for each orientation.

#### Data analysis

Automated cortical and subcortical segmentation of the T1-weighted and T2-weighted brain data was performed in Freesurfer (v6.0) (Fischl et al., 2002, 2004). For each hemisphere, we obtained the volume of the hippocampus (HC) and entorhinal cortex (EC) in the MTL for our main analysis. We also obtained striatal volumes, including left and right globus pallidus, putamen, caudate and accumbens for exploratory analyses (see *Supplemental Information*). Segmentations of these areas were checked visually and manually adjusted if necessary.

In addition to the Freesurfer segmentations, we obtained separate volumetric measures of the anterior and posterior hippocampus in each hemisphere. The ultra-high-resolution T2-weighted 0.5mm isotropic medial temporal lobe scans were submitted to automated segmentation using HIPS, an algorithm previously validated to human raters specialized in segmenting detailed neuroanatomical scans of the hippocampus (Romero et al., 2017). Three independent raters were trained on segmenting the hippocampus at the uncal apex into aHC and pHC segments, and achieved a Dice coefficient of absolute agreement of 80%. Two of these raters independently segmented all participants using the 0.5 mm T1-weighted scans. The T2-weighted medial temporal lobe scans were registered to the T2-weighted whole-brain scans, which were in turn registered to the T1-weighted whole-brain scans, and the combined transform was used to place the rater landmarks on the detailed medial temporal lobe scans. Finally, the total number of voxels in each subregion was multiplied by the volume of each voxel to obtain a total aHC and pHC volume.

To account for differences in head size, all regional volumes were corrected for total intracranial (IC) volume obtained from Freesurfer. This was done by first estimating the slope *b* of the regression line of each regional volume on the IC volume across the 33 participants included in the analysis. Next, each regional volume was adjusted for the IC volume as: adjusted volume = raw volume - *b* ⨉ (IC volume - mean IC volume).

### Motor learning tasks

#### General procedure

Thirty-four participants performed an error-based and a reward-based motor learning task. We attempted to fully counterbalance the tasks across participants; The first 19 participants performed the error-based motor learning task before performing the reward-based motor learning task, with the next 15 participants performing the reward-based motor learning task before the error-based motor learning task. The reward-based task took about 25 minutes to complete and the error-based task took about 65 minutes to complete.

#### Setup

Participants were seated at a table, with their chin and forehead supported by a headrest placed ~50 cm in front of a vertical LCD monitor (display size 47.5 × 26.5 cm; resolution 1920 × 1080 pixels) on which the stimuli were presented (Fig. 1A). Participants performed reaching movements by sliding a stylus across a digital drawing tablet (active area 311 × 216 mm; Wacom Intuous) placed on the table in front of the participant. Movement trajectories were sampled at 100 Hz by the digitizing tablet. Vision of the hand and tablet was occluded by a piece of black cardboard attached to the headrest. In the error-based learning task, eye movements were tracked at 500 Hz using a video-based eye tracker (Eyelink 1000; SR Research) placed beneath the monitor. The eye movement data were not analyzed in this study. The stimuli and motor learning tasks are described in detail below.

**Figure 1.**
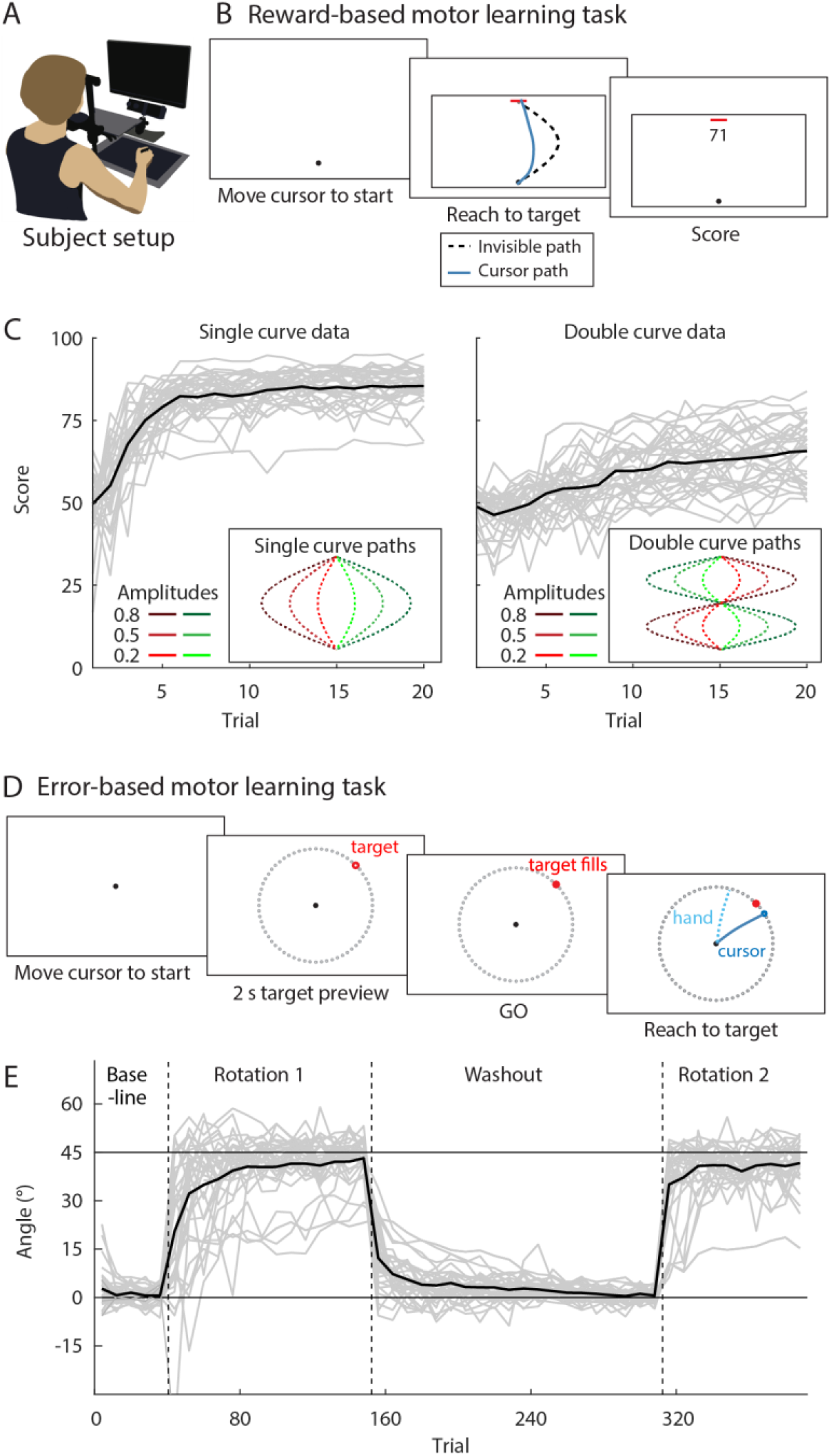
Experimental tasks and learning curves averaged across participants. (A) Setup. Participants made reaching movements by sliding a pen across a digitizing tablet without vision of the hand. The stimuli and a cursor representing the hand position were presented on a monitor. (B) Reward-based motor learning task. Participants were instructed to ‘copy’ an invisible path (dashed black line), with a score between 0 and 100 indicating how close their drawn path (blue line) was to the hidden path. (C) Learning across trials for single (left panel) and double invisible paths (right panel). Each grey line is an individual participant (n=33), with the black line representing the average across participants. Insets depict the six single curve invisible paths and the six double curve invisible paths used in the task. (D) Error-based motor learning task. Participants made center-out reaching movements to visual targets (red dot) on a ring of landmarks (small grey dots) with veridical cursor feedback (not shown) or under a 45° rotation of the cursor feedback (blue line; hand direction is shown in light blue). (E) Learning curves (left panel) across the baseline, rotation 1, washout, and rotation 2 block. Each grey line represents an individual participant (n=33), the black line represents the mean across participants.

#### Reward-based motor learning

##### Task

Our task was inspired by the reward-based learning task designed by Dam and colleagues (Dam et al., 2013). Participants performed reaching movements from a start position to a target line by sliding the stylus across the tablet. They were instructed to “find an invisible curved path by drawing paths on the tablet and evaluating your score for each attempt”. Participants started with a practice block of 10 trials, in which they traced a visible, straight line between the start position and the target, to become familiar with the task and the timing requirement of performing the movement within 2 s. Next, participants performed 12 blocks, each containing 20 attempts to copy an invisible path, which differed in each block.

Each trial started with the presentation of a start position (5 mm radius circle; Fig. 1B). After the participant had moved the cursor to the start position and held it there for 200 ms, a horizontal target line (30 × 1 mm) would appear 15 cm in front of the start position, and a rectangular outlined box (320 × 170 mm) would appear around the start position and target. Next, participants drew a path from the start position to the target line while remaining in the box. After crossing the target line, the cursor disappeared, and a score between 0 and 100 was displayed centrally (for 1 s), indicating how close they were to the invisible path. Following this, all stimuli disappeared, and a new trial would start with the presentation of the start position and the reappearance of the cursor. If the movement duration was longer than 2 s, the score was not presented and the trial was repeated.

The invisible paths consisted of single curves (i.e., half sine waves; 6 blocks) and double curves (i.e., full sine waves; 6 blocks) of different amplitudes (± 0.2, 0.5 and 0.8 times the target distance; see inset of Fig. 1C), drawn between the start position and the center of the target line. Participants were not informed about the possible shapes of the invisible lines. The trial score was computed by taking the x position of the cursor at every cm travelled in the y-direction (i.e., 1, 2, 3, … and 15 cm), and computing the absolute difference in x position between the cursor and the invisible line at the corresponding y-distance. The sum of these errors was then normalized by dividing it by the sum of distances between a straight line and a curve with an amplitude of 0.5 times the target distance, and multiplied by 100 to obtain a score between 0 and 100 (negative scores were presented as 0).

All participants performed one practice block and 12 experimental blocks of trials. Ten different randomized orders of experimental blocks were created. Participant 1, 11, 21, and 31 performed the first order, participant 2, 12, 22, and 32 performed the second order, etc.

##### Data analysis

The median score in trials 11 to 20 of each block of 20 attempts were used as a measure of learning performance. We did not use trials 1-10 in our analysis based on our frequent observation that participants who learned fairly quickly often used exploratory strategies when encountering a new path, which often resulted in scores of, or around, zero on several trials (Fig. 2A provides a good example of such a participant). For each participant, we averaged the median scores across all single curves and across all double curves with the same amplitude. This resulted in two scores per participant.

**Figure 2.**
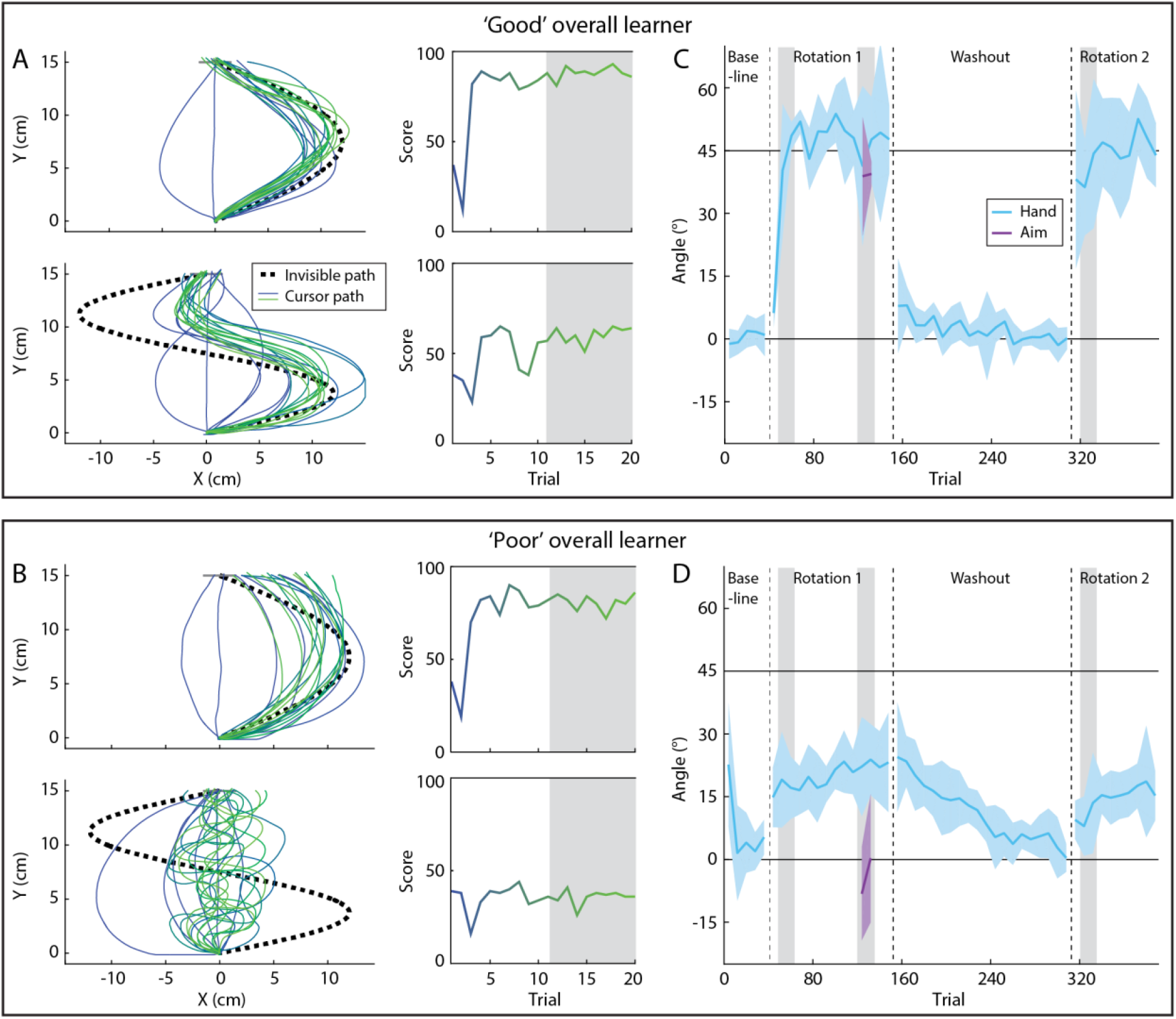
Example data of a ‘good learner’ and a ‘poor learner’. (A,C) Data of an example ‘good learner’. (A) Hidden path (dashed black line), drawn paths (blue and green lines), and score (blue to green gradient) for two blocks in the reward-based learning task. The median score in the last 10 trials of each block (grey shaded area) was used in further analyses. (C) Hand angle (light blue) and reported aiming angle (purple) relative to the target angle during the error-based learning task. Each data point represents the median of a set of eight trials, and the shading represents ± one standard deviation. The mean scores across sets 2 and 3 (early learning) of the rotation blocks were used in further analyses, as well as the averaged aiming angle (grey shaded areas). In the baseline and washout blocks, a hand angle of zero would result in a target hit, and in the rotation blocks, a hand angle of 45° results in perfect compensation of the rotated cursor path, and thus a target hit. (B,D) Same as (A,C), but for a ‘poor learner’.

#### Error-based motor learning

##### Task

Participants performed center-out reaching movements from a start position to one of eight visual targets presented on a 10 cm radius ring around the start position. Participants were instructed to hit the target with their cursor by making a fast reaching movement on the tablet, ‘slicing’ through the target. The ratio between movement of the tip of the stylus and movement of the cursor presented on the screen was 1:2, so that a movement of 5 cm on the tablet corresponded to a movement of 10 cm of the cursor. Participants first performed a baseline block in which they received veridical feedback about the position of the tip of the stylus, shown as a cursor on the screen. After performing a baseline block, participants performed a visuomotor rotation task, a task that has been used extensively to assess error-based learning (e.g., Cunningham, 1989; Krakauer et al., 2005). In this task, the movement of the cursor representing the hand position is rotated about the hand start location, in this experiment by 45° clockwise, requiring that a counterclockwise adjustment of movement direction be learned.

Each trial started with the participant moving the stylus to a central start position (5 mm radius circle; Fig. 1D). When the (unseen) cursor was within 5 cm of the start position, a ring was presented around the start position to indicate the distance of the cursor, so that the participant had to reduce the size of the ring to move to the start position. The cursor (4 mm radius circle) appeared when the cursor ‘touched’ the start position (9 mm distance). After the cursor was held within the start position for 500 ms, the target (6 mm radius open circle) was presented on an (imaginary) 10 cm radius ring around the start position at one of eight locations, separated by 45° (i.e., 0, 45, 90, 135, 180, 225, 270 and 315°). In addition, 64 non-target ‘landmarks’ (3 mm radius outlined circles, spaced 5.625° apart) were presented, forming a 10 cm radius ring around the start position. After a 2 s delay, the target would ‘fill in’ (i.e., color red), providing the cue for the participant to perform a fast movement to the target. If the participant started the movement before the cue, or more than 1 s after the cue, the trial was aborted and a feedback message indicating “Too early” or “Too late” appeared on the screen, respectively. In correctly timed trials, the cursor was visible during the movement to the ring and then became stationary for 1 s when it reached the ring, providing the participant with visual feedback of their endpoint error. When any part of the cursor overlapped with any part of the target, the target would color green to indicate a hit. If the duration of the movement was longer than 300 ms, a feedback message “Too slow” would appear on the screen.

In trials in the rotation block, the movement of the cursor was rotated by 45° clockwise around the start position. To assess the contribution of the explicit process of learning, participants performed several ‘reporting’ trials. These trials were performed at the end of the first rotation block to ensure that participants’ learning behavior would not be influenced, as the reporting procedure itself can increase the proportion of participants that implement a cognitive strategy (3). In reporting trials, participants were instructed to, before each reach movement, report the aiming direction of their hand for the cursor to hit the target. They did this by turning a knob with their left hand, to rotate a line on the screen, positioned between the start position and the ring, to align it with their strategic aimpoint. When satisfied with the direction of the line, the participant clicked a button positioned next to the knob, and the line disappeared. After a 1 s delay, the target filled in as a cue to execute the reach.

All participants performed 4 blocks of trials in total, where within each block, target locations were randomized within sets of eight: (1) A baseline block (5 sets of 8 trials; 40 in total), (2) a rotation block (10 sets of 8 trials without report + 2 sets of 8 reporting trials + 2 sets of 8 trials without report; 112 trials in total), (3) a washout block in which veridical cursor feedback was restored (10 + 10 sets with a 30 s break in between; 160 trials in total) and (4) a second rotation block to assess participants’ rates of re-learning (10 sets of 8 trials; 80 in total). Ten different randomized trial orders were created for the full experiment. Participant 1, 11, 21, and 31 performed the first order, participant 2, 12, 22, and 32 performed the second order, etc.

##### Data analysis

Trials in which the movement was initiated too early or too late (as detected online; 4% of trials) or in which the movement duration was longer than 300 ms (4% of trials), were discarded from the analysis. The median endpoint hand angle (i.e., the difference in angle between the target and the hand when the cursor crossed the ring) per set of eight trials was used as a measure of learning performance. To capture individual differences in the rate of early learning that correspond to the implementation of aiming strategies (Taylor et al., 2014; de Brouwer et al., 2018), we computed, for each participant, early learning scores in the first and second rotation block. To do this, we averaged the median in sets 2 and 3 of each of these blocks (excluding the first set in which participants often showed highly variable behavior). To derive a direct measure of the magnitude of the explicit component of learning, we used the average of the median reported aiming angle with respect to the target, obtained in the reporting trials (sets 11 and 12 in the rotation block). This resulted in three measures per participant.

### Relating learning measures and neuroanatomy

#### Data and statistical analysis

As a first exploratory step to determine relationships in subject performance across the error-based and reward-based learning tasks, we calculated Pearson correlations between the five learning scores across participants. Having identified patterns of covariation in subject-level performance across the two tasks, for our main analysis, we submitted the learning scores to a principal component analysis (PCA). This approach has three important advantages: (1) it identifies the main patterns of covariation both within and between tasks, (2) it reduces the number of behavioural variables to be used in further analyses, and (3) it provides us with uncorrelated measures of learning performance (i.e, principal components), suitable to use in linear regression. To do the PCA, we first transformed the variables from the error-based learning task, whereby all angles were converted to errors with respect to the target, such that zero corresponds to a target hit, negative errors (i.e., between −45° and 0°) correspond to no or partial compensation of the rotation, and positive errors correspond to overcompensation of the rotation. This transformation ensured that higher values on both the error-based and reward-based motor learning tasks were associated with better learning performance. We then standardized all scores before submitting them to the PCA. The principal components (PCs) were obtained using the pca function in Matlab, which uses a singular value decomposition algorithm to find PCs that capture the maximal variance in the data.

To test the hypothesis that better performance in the motor learning tasks is related to greater volumes of brain areas in the medial temporal lobe, we performed multiple linear regression analyses. All models were estimated using the fitlm function in Matlab, which returns a least-squares fit of the scores to the data. Our primary analysis included the left and right HC and EC volumes. To control for a potential effect of overall head size on learning performance, we also included each participant’s total intracranial volume, making a total of five neuroanatomical measures. For the first and second PC, we fitted a multiple linear regression model with the PC as the dependent variable, and the set of four regional volumes plus the IC volume as predictors. Previous studies have reported differential relationships between the anterior and posterior parts of the hippocampus and memory (e.g., Maguire et al., 2000). Therefore, we performed a secondary analysis, including the left and right anterior and posterior HC volume as predictors, and the IC volume as a confounder.

## Results

In order to determine the relationship between motor performance in reward-based and error-based learning tasks, and the extent to which the size of hippocampal and entorhinal cortex may be associated to such learning, we collected high-resolution structural MRI scans from participants (N=34) prior to performing two separate motor learning tasks outside the scanner. In the reward-based learning task, participants learned to copy an invisible, curved path through trial and error, using only a score (between 0 and 100 points) to improve their performance. This score, presented at the end of each trial, indicated how closely the participants’ drawn path corresponded to the invisible path (Fig. 1B). Participants drew these paths on a digital drawing tablet from a start to a target position displayed on a vertical monitor (Fig. 1A), and were instructed to maximize their score. To obtain a representative measure of each participant’s reward-based learning rate and ability, we had participants perform this task for 12 different invisible paths, with 20 attempts for each. Participants were naive to the possible shapes of the paths, which were shaped as single curves (i.e., half sine waves) and double curves (i.e., full sine waves) between the start and target position, with different amplitudes (see Fig. 1C). Because participants received only visual feedback about their path trajectory—and never the rewarded path—they did not receive error-based information that could be used to guide learning. By design, this reward-based task requires implementing a search strategy to first find the invisible path and then refine the drawn path, and we thus predict that participants who perform well in this task are better at implementing such strategies.

For the error-based learning task, we used the classic visuomotor rotation learning paradigm (Cunningham, 1989), wherein participants had to adjust their movements to a 45° rotation of the cursor movement, which represented participants’ hand movements, in order to hit visual targets (Fig. 1D). Participants performed center-out reaching movements on the drawing tablet to one of eight targets displayed on a monitor. After a baseline phase with veridical cursor feedback, participants were exposed to the 45° visuomotor rotation of the movement of the cursor, requiring an adjustment of the reaching movement in the opposite direction. Learning in this task consists of two components: automatic, implicit adjustments of the reach direction, resulting in gradual changes in performance, and the implementation of an aiming strategy to counteract the rotation, resulting in fast changes in performance (Redding and Wallace, 1993; Taylor et al., 2014). Our previous work has shown (de Brouwer et al., 2018) — and we predict here — that faster and more complete learning is largely driven by the use of an aiming strategy, used to counteract the rotation. At the end of the first block of rotation trials, we assessed this aiming strategy by asking participants to report the intended aiming angle by turning a knob with their left hand to rotate a line on the screen at the start of the trial, before executing the reach (Taylor et al., 2014). Learning was then ‘washed out’ by restoring veridical cursor feedback, after which the visuomotor rotation was re-instantiated to assess the rate of re-learning.

### Performance in reward-based and error-based motor learning is related

The black traces in Figure 1C and 1E show the learning curves, averaged across all participants, for the reward-based and error-based learning tasks, respectively. These figures demonstrate that participants learned to increase their scores in the reward-based task and change their hand angle in the error-based task across trials. However, these group-averaged results may be somewhat misleading, as they obscure significant intersubject variability in both the rates and levels of learning obtained (see gray traces in Fig. 1C,E, which depict single participants). For example, Figure 2 shows the behavior of two participants, one ‘good’ overall learner and one ‘poor’ overall learner, in both the reward-based learning task and the error-based learning task. Figure 2A and 2B depict the paths that the participants drew (left panel) and the corresponding scores (right panel), in two blocks of the reward-based learning task for a single (top) and double curve (bottom) with the largest amplitude (blocks 4 and 11 for the participant in Fig. 2A; blocks 11 and 10 for the participant in Fig. 2B). While both participants quickly converged on a good solution for the single curve, resulting in scores close to 100, the movements of the participant in Figure 2A resemble the invisible curve more closely. In addition, while the participant in Figure 2A quickly converges upon a solution that has a similar shape to the invisible double curve, the participant in Figure 2B never learns to draw that same double curve, and their score remains low.

Figure 2C and 2D show, for the same two participants, the median hand angle (in blue) for each bin of eight trials across the error-based learning task, as well as the reported aiming angle (in purple) assessed near the end of the first rotation block. Appropriate corrections for the visuomotor rotation are plotted as positive values; that is, a hand angle of 45° corresponds to full compensation for the rotation. The participant in Figure 2C shows quick adjustment of the hand angle towards 45° in the first and second rotation block, and a quick return towards 0° in the washout block. Such fast learning is associated with a large contribution of an aiming strategy, consistent with their reported aiming angles around 39°. The participant in Figure 2D, by contrast, shows only gradual adjustments of the hand angle in the rotation and washout blocks, and correspondingly reports aiming values around 0°, suggesting that learning in this participant is mainly driven by the implicit process. Overall, the participant in Figure 2A,C showed better learning performance in both tasks than the participant in Figure 2B,D.

For each participant, we obtained two learning scores for the reward-based learning task (single and double curves) and three learning scores for the error-based learning task (early and late learning in rotation block 1 and 2, and the reported aiming angle; Fig 3A). Across the entire group of participants, we observed several significant correlations in the learning scores both within and between the two tasks (Fig. 3B). Notably, the latter demonstrates clear patterns of covariation in subject-level performance across both the error-based and reward-based motor learning tasks. To derive single participant measures of learning that capture these patterns of covariation, and that can be used to relate overall learning performance to the neuroanatomical data collected in these same participants, we performed a principal component analysis (PCA) on the learning scores (see *Methods* for details). We found that the first (PC1) and second principal components (PC2) explained 53.8% and 22.3% of the variance in the data (76.1% overall), respectively. The projection plots in Figure 3C (left panel) allows for a straightforward interpretation of these PCs, directly showing both the magnitude and sign of the loading of each of our 5 learning measures onto PC1 and PC2. Notably, PC1 has positive loadings for all of the learning measures, indicating that this single component captures overall learning performance. Indeed, PC1 shows significant positive correlations with *all* behavioral learning measures from both tasks (ranging from *r*=0.55 to *r*=0.82, all *p*<0.001, see Fig. 3C, right panel). In other words, PC1 provides a single scalar measure that distinguishes between relatively ‘good’ versus ‘poor’ learning performance across *both* the reward-based and error-based learning tasks. The second principal component (PC2) broadly distinguishes between performance in the reward-based and error-based learning task, with positive loadings for the reward-based learning scores and negative loadings for learning in the rotation blocks. However, PC2 explains a relatively small portion of the overall behavioral variance (22.3%), limiting its interpretational value and its use in further analyses. Taken together, our dimensionality-reduction approach on the behavioral learning data demonstrates that subject-level performance in both tasks is highly related, as a single latent variable (PC1) captures whether participants are good learners in both the reward-based learning and in the error-based learning task.

**Figure 3.**
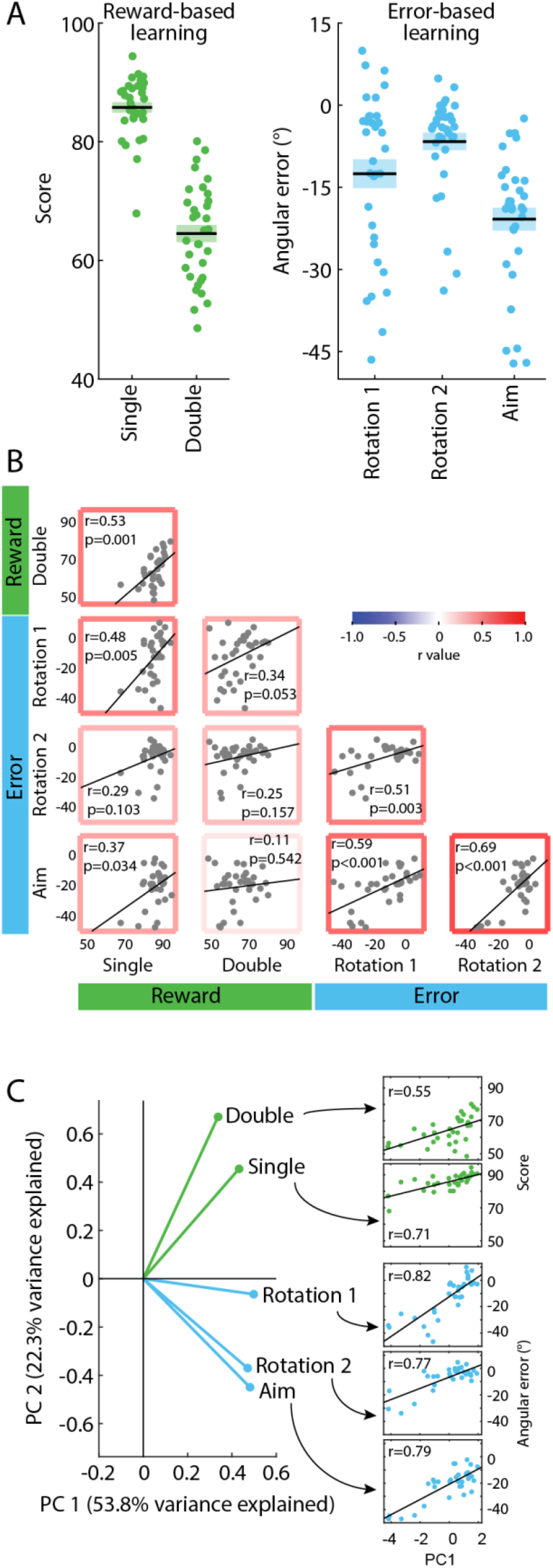
Learning performance in the error-based and reward-based tasks is related and is captured by a single latent variable. (A) Distribution of scores for single and double curve performance in the reward-based learning task, and distribution of angular errors during early learning in the first and second rotation block, and reported aiming errors. Each dot indicates the mean value of one participant, the horizontal line indicates the mean across participants, and the shaded area indicates the standard error of the mean. (B) Scatterplots and associated Pearson correlations (uncorrected) within and between scores in the reward-based and error-based motor learning tasks across participants (n=33). This demonstrates subject-level covariation in learning performance both within and between the two tasks. (C) Principal component analysis loadings for the first and second principal component (PC) at left, and Pearson correlations between the first principal component and learning scores at right. This shows that PC1 provides a useful proxy for learning performance across both tasks. In (B) and (C), each dot represents one participant, and the black line represents the best fit regression line.

### Larger entorhinal volume is associated with better error- and reward-based motor learning

Having clearly established that subject-level performance in reward-based and error-based learning is related and that this pattern of covariation can be captured by a single measure (i.e., PC1), our next aim was to determine whether this variation in performance is associated with the neuroanatomy of the MTL. To this end, we performed multiple linear regression analyses using right and left hippocampus (HC) and entorhinal cortex (EC) volumes as predictors and PC1 as the outcome variable (Fig. 4AB), corrected for total intracranial volume (see *Methods*). We also included total intracranial (IC) volume in our model to account for a potential effect of overall head size. Figure 4C shows the standardized regression coefficients and 95% confidence interval of the regression models for each PC. For PC1 — our measure of good vs. poor performance in both tasks — the model significantly explained the variance in PC1 score (model *F*(5)=4.050, *p*=0.007; *R*_*2*_=42.9%), with right EC volume being a significant predictor (*t*=3.689, *p*=0.001; see Figure 4-1). That is, larger right entorhinal volume corresponded with higher scores on PC1, or better overall learning in both tasks. Notably, we found no significant predictors of PC2 score (model *F*(5)=0.410, *p*=0.806, *R*_*2*_=7.1%), the measure that broadly distinguished between performance in the reward-based and error-based learning task. This lack of effect might not be surprising given that the percentage of variance in our motor learning data that was explained by the second principal component was fairly small (22.3%).

**Figure 4.**
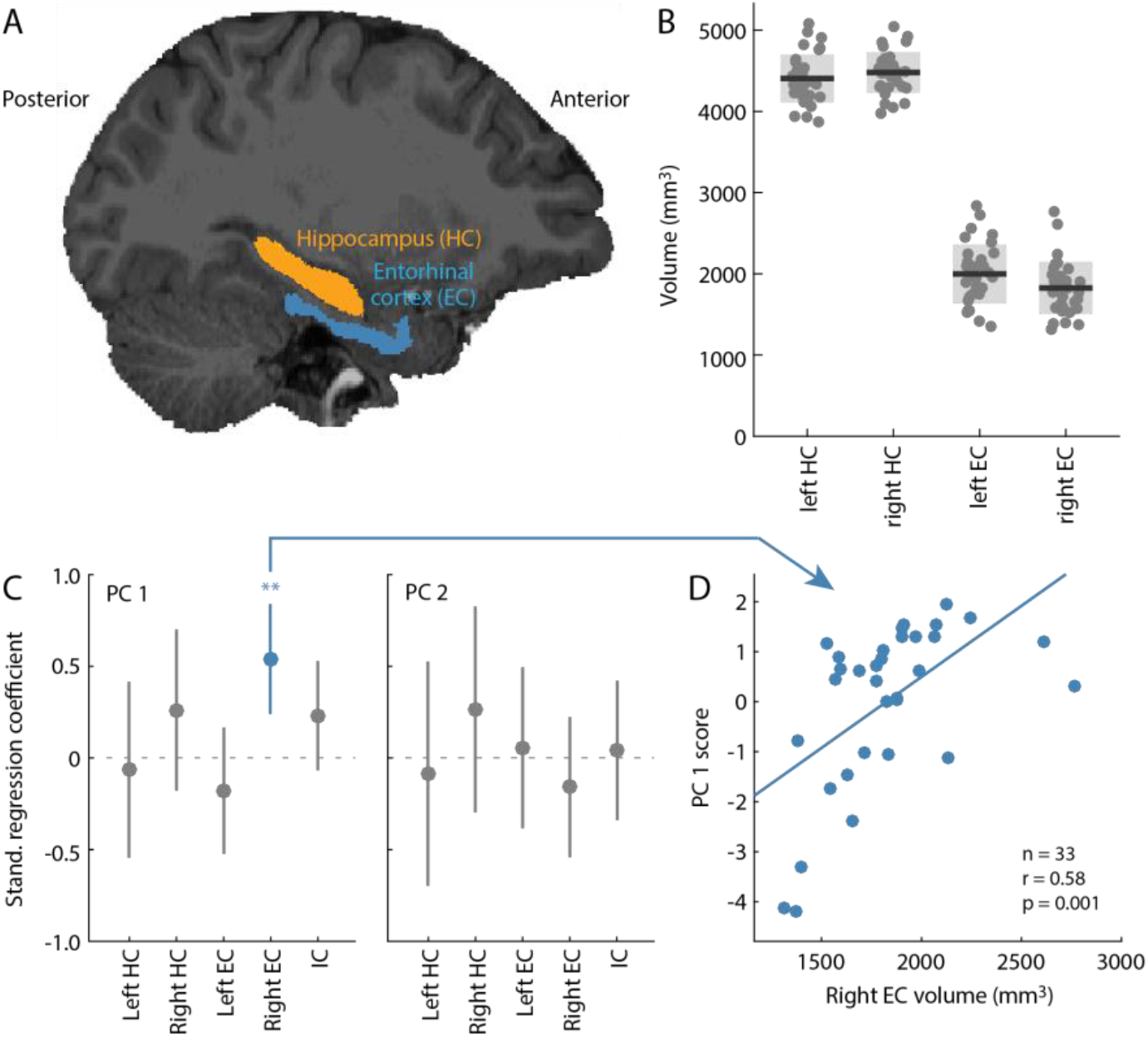
Larger entorhinal volumes are uniquely associated with better overall motor learning. (A) Illustration of the segmented hippocampus (orange) and entorhinal cortex (blue) in an example participant. (B) Volume of the left (L) and right (R) hippocampus (HC) and entorhinal cortex (EC), corrected for total intracranial volume (see *Methods*). Each dot depicts an individual participant (n=33), the dark grey line indicates the mean across participants, and the light grey area indicates the standard deviation. (C) Standardized regression coefficients and their corresponding 95% confidence intervals of the regression models to predict principal component 1 (PC 1; left panel) and principal component 2 (PC 2; right panel) based on the left and right hippocampus and entorhinal volumes, and the total intracranial volume (IC). See Figure 4-1 and 4-2 for all model coefficients and significance values. (D) Individual partial correlation between right entorhinal volume and PC 1 score. The line represents the best fit regression line.

As a secondary analysis, we performed a linear regression with the anterior and posterior hippocampus as separate predictors, as previous studies have reported differential relationships between these individual parts of the hippocampus and memory (e.g., Maguire et al., 2000). However, here we again did not find significant relationships between left and right aHC and pHC volume and the score on PC1 (model *F*(5)=0.360, *p*=0.871, *R2*=6.2%) or PC2 (*F*(5)=0.440, *p*=0.817, *R*_*2*_=7.5%; Figure 4-2).

Taken together, the results of these regression analyses indicate that better performance in both reward-based and error-based learning is associated with larger right entorhinal volume.

## Discussion

While previous work in motor learning has often studied error-based and reward-based learning processes in isolation from one another, recently there has been increased interest in understanding how these separate learning processes interact at the behavioral and neural levels. Here we find a strong relationship in intersubject variability between error-based and reward-based motor learning, showing that learning performance across tasks is correlated and can be explained by a single, latent variable. Our measures of learning and the nature of the tasks used suggest that this latent variable captures participants’ use of cognitive strategies during learning, with higher scores on this variable being associated with faster and better overall learning in both tasks. We further show, using structural neuroimaging and regression analyses with participants’ hippocampus and entorhinal cortical volumes as predictors, that higher scores on this latent variable, and thus faster and better overall learning, is associated with larger right entorhinal cortex volumes. Together, these findings suggest that a shared strategic process underlies individual differences in error-based and reward-based motor learning, and that this process is associated with structural differences in entorhinal cortex.

Considerable computational and neural work has argued for a division of labor between the neural circuits that support error-based and reward-based learning (Doya, 1999, 2000; Daw and Doya, 2006; Shadmehr and Krakauer, 2008; Ito and Doya, 2011; Makino et al., 2016). According to this prevailing view, cortico-cerebellar pathways are responsible for error-based learning whereas cortico-striatal pathways are responsible for reward-based learning. Such distinctions, however, have often been reliant on indirect comparisons between different studies, and have been influenced by sampling biases in neural recording sites across different tasks. For instance, conventional views on error-based learning have suggested that adaptation is a primarily automatic mechanism, immune to reward-based feedback (Doya, 2000; Shadmehr and Krakauer, 2008). However, more recent behavioral evidence suggests that these two learning processes, while separable (Izawa and Shadmehr, 2011; Cashaback et al., 2017), interact during sensorimotor learning (Shmuelof et al., 2012; Taylor and Ivry, 2014; Galea et al., 2015; Nikooyan and Ahmed, 2015). Such interactions are likely to be supported by the recent demonstration of direct anatomical connections between the cerebellum and striatum (Bostan and Strick, 2018). These bidirectional connections could explain recent neural findings from rodents showing that the cerebellum, besides processing direction-related errors, also represents various aspects of reward-related information during task performance (Wagner et al., 2017; Heffley et al., 2018; Kostadinov et al., 2019; Larry et al., 2019). Together, this emerging evidence suggests that error-based and reward-based learning processes are closely intertwined at both the behavioral and neural levels.

There is also emerging evidence to suggest that both error-based and reward-based processes are mediated through the use of cognitive strategies implemented during learning. In error-based adaptation, the contribution of this explicit, declarative process to learning has been well-established behaviorally (Redding and Wallace, 1993; Fernandez-Ruiz et al., 2011; Taylor and Ivry, 2011; Taylor et al., 2014; Bond and Taylor, 2015; Haith et al., 2015; de Brouwer et al., 2018). Recent evidence from our group further indicates that faster learning across participants is linked to individual differences in the magnitude of the cognitive strategy (de Brouwer et al., 2018), which drives rapid changes early in the learning process. In reward-based learning, by contrast, the contribution of cognitive strategies to performance have received comparably little attention, and is only beginning to be established. As one example, recent work, wherein participants were only provided with reward-based feedback (binary success/failure) to perform a visuomotor rotation task, has shown that good versus poor learning is related to the implementation of a cognitive component (Holland et al., 2018). This was evidenced by the observed reduction in reach angle when participants were required to remove their aiming strategy (see also Codol et al., 2018). It was also evidenced by the observation that the reward-based learning was impaired when (1) participants had to perform a dual task (a separate mental rotation task) that divided their cognitive load (Holland et al., 2018), or when (2) participants’ reaction times were constrained (Codol et al., 2018), such that they could not implement the strategy (Haith et al., 2015). To date, work examining the link between error- and reward-based learning has focused on how reinforcement signals (e.g., binary success/failure) shape learning in traditionally error-based tasks (Shmuelof et al., 2012; Galea et al., 2015; Cashaback et al., 2017). By contrast, our current behavioral findings show that, even when reward- and error-based learning is studied separately (and in very different tasks), learning performance in both tasks is highly related — so much so that a single latent variable can explain a significant proportion of intersubject variability in performance across both types of learning.

Another novel result in our study was our finding that a larger right entorhinal volume was associated with better overall learning in both the reward-based and error-based motor learning tasks. The entorhinal cortex has been shown to support a wide range of cognitive functions that would have bearing on various features of our motor learning tasks. Classically, the entorhinal cortex, together with neighboring areas in the medial temporal lobe, has been implicated in spatial navigation and memory through electrophysiological studies in rodents. These studies showed that place cells in the hippocampus (O’Keefe and Dostrovsky, 1971) and grid cells in the entorhinal cortex (Hafting et al., 2005) form a map-like representation of the environment. Grid cells have also been demonstrated in primate entorhinal cortex, even in the absence of locomotion, when the animal is simply exploring a visual scene with its eyes (Killian et al., 2012, 2015). Such observations have recently been extended to humans with functional MRI (Julian et al., 2018; Nau et al., 2018), and there is even evidence suggesting that mere shifts in covert attention (i.e., in the absence of overt eye movements), also elicits grid-cell-like responses in the entorhinal cortex (Wilming et al., 2018). Together, these and other findings (Bellmund et al., 2016; Constantinescu et al., 2016; Horner et al., 2016) have begun to reshape our understanding of the role of the entorhinal cortex in visual-spatial memory, and in cognitive operations more generally. An influential hypothesis is that the hippocampal-entorhinal system supports a cognitive map, an idea that was originally proposed to explain findings in rodents (Tolman, 1948; O’Keefe and Nadel, 1978) and later extended to humans (for review see Epstein et al., 2017). This hypothesis proposes that the brain creates flexible representations of the environment to not only support memory but also guide future decisions and effective (motor) behavior (Schiller et al., 2015; Garvert et al., 2017; Bellmund et al., 2018).

In the context of the current study, we expect cognitive and spatial maps to be utilized during the exploration of visuomotor space in our curve drawing (reward-based) and visuomotor rotation (error-based) tasks. Studies using fMRI in healthy adults, and neural recordings or electrical stimulation in pre-surgical patients, have provided evidence that the entorhinal cortex supports the encoding of goal direction and distance, relative locations, and the clockwise or counterclockwise direction of routes (Jacobs et al., 2010, 2016; Miller et al., 2013, 2015; Chadwick et al., 2015; Goyal et al., 2018; Qasim et al., 2019). While our motor learning tasks did not involve navigation in VR, the encoding of goal directions (in the visuomotor rotation task) and trajectories to the goal (in the curve drawing task) were critical to learning. If the entorhinal cortex is important for representing such spatial information, then its size may predict the ability to perform tasks — perceptual and motor — that recruit such representations. Studies investigating the relation between neuroanatomy and performance have associated greater gray matter volume in the entorhinal cortex with better scene recognition (Whiteman et al., 2016), spatial memory (Hartley and Harlow, 2012), navigation to memorized object locations in VR (Sherrill et al., 2018), as well as the lifetime amount of video gaming (Kühn and Gallinat, 2014). Here, we extend these general observations to include the previously unexplored domain of motor learning, showing an association between right entorhinal volume and overall performance in error-based and reward-based learning tasks. Given that motor learning has a strong visual-spatial component (particularly so in our tasks), we find it noteworthy that it is the right, and not left, entorhinal cortex that is associated with the processing and integration of visual-spatial information (Dalton et al., 2016).

## Author contributions

Conceptualization and methodology AJdB, JP, JPG, JRF; investigation AJdB; software AJdB; formal analysis AJdB, JP, MRR; visualization AJdB, JPG, MRR; writing - original draft AJdB, JPG; writing - review and editing AJdB, JP, JPG, JRF, MRR; supervision AJdB, JP, JPG; project administration AJdB, JP, JPG; resources JP, JPG, JRF; funding acquisition JP, JPG, JRF.

## Acknowledgements

This work was supported by operating grants from the Canadian Institutes of Health Research (CIHR) awarded to J.R.F. and J.P.G. (MOP126158). J.P.G. and J.P. were supported by Natural Sciences and Engineering Research Council (NSERC) Discovery Grants, as well as funding from the Canadian Foundation for Innovation. The authors thank Mohammed Albaghdadi, Olivia Broda, Sydney Dore, Kate McKenzie and Reem Toubache for help with data collection, and Martin York for technical support.

