## Extended Data for "Variability in error-based and reward-based human motor learning is associated with entorhinal volume"

**Figure 4-1.** Contributions of hippocampus and entorhinal volume to PC1 and PC2 score

|  | Variable | B | SE | 95% CI | | Beta | SE | 95% CI | | t | p | Model F | Model df | Model p | Model R^2^ | Adj. R^2^ |
| --- | --- | --- | --- | --- | --- | --- | --- | --- | --- | --- | --- | --- | --- | --- | --- | --- |
|  |  |  |  | lower | upper |  |  | lower | upper |  |  |  |  |  |  |  |
| PC1 | L HC | 0.000 | 0.001 | -0.003 | 0.002 | -0.064 | 0.234 | -0.545 | 0.416 | -0.275 | 0.786 | 4.050 | 5 | 0.007 | 0.429 | 0.323 |
|  | R HC | 0.002 | 0.001 | -0.001 | 0.005 | 0.261 | 0.215 | -0.180 | 0.702 | 1.216 | 0.234 |  |  |  |  |  |
|  | L EC | -0.001 | 0.001 | -0.002 | 0.001 | -0.179 | 0.168 | -0.524 | 0.167 | -1.061 | 0.298 |  |  |  |  |  |
|  | R EC | 0.003 | 0.001 | 0.001 | 0.004 | 0.540 | 0.146 | 0.240 | 0.840 | 3.689 | 0.001 |  |  |  |  |  |
|  | IC | 0.000 | 0.000 | 0.000 | 0.000 | 0.231 | 0.145 | -0.067 | 0.530 | 1.588 | 0.124 |  |  |  |  |  |
| PC2 | L HC | 0.000 | 0.001 | -0.002 | 0.002 | -0.086 | 0.299 | -0.699 | 0.526 | -0.289 | 0.775 | 0.410 | 5 | 0.837 | 0.071 | -0.102 |
|  | R HC | 0.001 | 0.001 | -0.001 | 0.003 | 0.265 | 0.274 | -0.298 | 0.827 | 0.966 | 0.343 |  |  |  |  |  |
|  | L EC | 0.000 | 0.001 | -0.001 | 0.001 | 0.055 | 0.215 | -0.385 | 0.496 | 0.257 | 0.799 |  |  |  |  |  |
|  | R EC | -0.001 | 0.001 | -0.002 | 0.001 | -0.159 | 0.187 | -0.542 | 0.224 | -0.852 | 0.402 |  |  |  |  |  |
|  | IC | 0.000 | 0.000 | 0.000 | 0.000 | 0.041 | 0.186 | -0.340 | 0.422 | 0.222 | 0.826 |  |  |  |  |  |

L = left, R = right, HC = hippocampus, EC = entorhinal cortex, IC = intracranial

**Figure 4-2.** Contributions of anterior and posterior hippocampus volume to PC1 and PC2 score

|  | Variable | B | SE | 95% CI | | Beta | SE | 95% CI | | t | p | Model F | Model df | Model p | Model R^2^ | Adj. R^2^ |
| --- | --- | --- | --- | --- | --- | --- | --- | --- | --- | --- | --- | --- | --- | --- | --- | --- |
|  |  |  |  | lower | upper |  |  | lower | upper |  |  |  |  |  |  |  |
| PC1 | L aHC | 0.000 | 0.003 | -0.007 | 0.007 | 0.024 | 0.301 | -0.593 | 0.642 | 0.081 | 0.936 | 0.360 | 5 | 0.871 | 0.062 | -0.111 |
|  | L pHC | 0.001 | 0.003 | -0.006 | 0.007 | 0.058 | 0.246 | -0.446 | 0.562 | 0.236 | 0.816 |  |  |  |  |  |
|  | R aHC | 0.000 | 0.003 | -0.007 | 0.007 | -0.023 | 0.329 | -0.699 | 0.653 | -0.070 | 0.945 |  |  |  |  |  |
|  | R pHC | 0.001 | 0.004 | -0.006 | 0.008 | 0.051 | 0.252 | -0.466 | 0.567 | 0.201 | 0.842 |  |  |  |  |  |
|  | IC | 0.000 | 0.000 | 0.000 | 0.000 | 0.231 | 0.186 | -0.151 | 0.613 | 1.240 | 0.226 |  |  |  |  |  |
| PC2 | L aHC | 0.003 | 0.002 | -0.002 | 0.007 | 0.365 | 0.299 | -0.248 | 0.978 | 1.222 | 0.232 | 0.440 | 5 | 0.817 | 0.075 | -0.096 |
|  | L pHC | 0.000 | 0.002 | -0.004 | 0.004 | -0.003 | 0.244 | -0.503 | 0.498 | -0.010 | 0.992 |  |  |  |  |  |
|  | R aHC | -0.002 | 0.002 | -0.006 | 0.003 | -0.246 | 0.327 | -0.917 | 0.425 | -0.751 | 0.459 |  |  |  |  |  |
|  | R pHC | -0.001 | 0.002 | -0.006 | 0.003 | -0.137 | 0.250 | -0.650 | 0.375 | -0.549 | 0.587 |  |  |  |  |  |
|  | IC | 0.000 | 0.000 | 0.000 | 0.000 | 0.041 | 0.185 | -0.339 | 0.421 | 0.222 | 0.826 |  |  |  |  |  |

L = left, R = right, aHC = anterior hippocampus, pHC = posterior hippocampus, IC = intracranial
